# Canadian science graduate stipends lie below the poverty line

**DOI:** 10.1101/2024.11.06.622240

**Authors:** Andrew Jeffrey Fraass, Thomas J Bailey, Kayona Karunakumar, Andrea E. Wishart

**Author notes:** These authors contributed equally to this work; Fraass & Bailey are joint first authors. Author Contributions: Project administration (TB), project design (AJF, TB, AEW), data and code management (TB, AJF), data collection (all authors), analysis (AJF, TB, AEW), writing (all authors) Thomas Bailey: Project administration, data management, data collection, analysis, writing Andy Fraass: Data collection, analysis, writing Kayona Karunakumar: Data collection, writing Andrea Wishart: Data collection, analysis, writing.

## Abstract

Despite the critical role of graduate students in the Canadian research ecosystem^1^, students report high levels of financial stress^2^. We collected graduate minimum stipends and tuition data from all university graduate programs in Canada in Ecological Sciences/Biology and Physics, along with cost of living measures for the cities in which they reside. This data is heterogeneous, complex, and in many cases simply not publicly available, making it challenging for potential graduate students to understand what support they should expect. We show Canadian minimum stipends are at values almost exclusively below the poverty threshold. Only two of 140 degree programs offered stipends which meet cost of living measures after subtracting tuition and fees. For graduate programs which offered a minimum guaranteed stipend, the average minimum domestic stipend is short ∼Can$9,468 (international ∼Can$16,899) of the poverty threshold after accounting for payment of tuition and fees. On average, approximately 34% of a minimum stipend is returned to the university by a domestic Canadian student and 78% (57% median) by an international student, though there are important caveats with the international student comparison. While international comparison is difficult, the highest Canadian minimum stipend is roughly equivalent or lower than the lowest stipend within the largest dataset of United States of America (US) Biology stipends ^3^, and lower than the United Kingdom (UK) stipend. University endowment correlates with minimum stipend amount but intra- and inter-institutional differences suggest it is not solely institutional wealth which improves graduate pay. Canada is behind comparable countries in funding the next generation of scientists. Canadians who desire higher STEM education have three options: hope for significantly higher guaranteed support from a supervisor, department, or awards; incur substantial debts; or emigrate.

## Introduction

Graduate students are a primary engine of scientific work for the academic system (Larivière 2012). Despite this critical role in the Canadian research ecosystem, students report high levels of financial stress^2^ exacerbated by increasing cost of living^4^.

The following is a general description of the financial model of Canadian graduate studies in natural science: stipends provide financial assistance to students for living expenses (e.g., tuition, rent), as opposed to financial support for the costs associated with the research itself (e.g., reagents, equipment)^5^. Graduate students in Canada pay tuition and fees to their institution; waivers are rarely provided. When a stipend is provided, tuition and fees are paid by the student to their institution using their stipend, which itself was previously paid by the institution to the student.

Departments or equivalent academic units frequently set minimum stipend levels but individual principal investigators (PIs) can choose to exceed that amount. Stipends are typically funded from multiple sources: the institution (e.g., faculty of graduate studies provides the department funds based on number of students in their programs), a PI’s research grants (see below), student awards, and/or teaching assistantships. TAships are a core part of the funding package, with the salary earned from this job often included in the value of the stipend.

The Natural Sciences and Engineering Research Council of Canada (NSERC) Discovery Grant (DG) is a 5-year operating grant awarded to individual PIs and the foundation of most academic Canadian natural science graduate funding and research. Grant applications are judged evenly across three categories: a record of past research, highly qualified personnel (HQP; students, postdocs, and other mentees) produced by the researcher, and a proposed project^6^. Quality mentorship is therefore paramount in acquiring funding. Policy requires NSERC panels to not judge a researcher based on the number of HQP^6^; rather, applicants are judged on mentorship quality, post-mentorship careers, and so on. However, an incentive still remains to have more students, as publications and citations correlate positively with the number of HQP^7^. This pressure to train a large number of students with the relatively small operating grants contributes to the suppression of stipend minimum support.

This study tests if the guaranteed minimum stipends for graduate students are a) enough to live on, b) are comparable between programs and fields, c) between universities and provinces, and d) against two peer nations (US and UK).

### Positionality

Given the impact of career stage and nationality on perspectives about graduate programs, we summarize our positionality thusly:

*Bailey* is a British PhD student (Physics) at the University of Ottawa (Canada). He currently receives stipend support though a NSERC grant awarded to his supervisor. He is a department steward for his TA union and is on the Executive Council of Support Our Science.

*Fraass* is an American tenure track Assistant Professor (Micropaleontology) at the University of Victoria (Canada). His graduate degrees and two postdoctoral positions were US based, followed by three years in England as a postdoc and fellow. He currently receives funding from NSERC and mentors graduate students.

*Karunakumar* is a Canadian early-career science policy professional. Her undergraduate and graduate education includes training in public policy and government relations. She is pursuing a career in science policy. Additionally, she serves as a volunteer for Evidence for Democracy.

*Wishart* is a Canadian early-career academic publishing professional currently employed by Canadian Science Publishing. Her degrees (BSc, MSc, PhD) in biology were at Canadian universities, save for one year abroad (UK). She previously held institutional scholarships and received stipend support through operating grants (e.g., NSERC DG) awarded to her supervisors. In addition to leadership roles advocating for graduate students and performing research on graduate student experiences, she co-founded Support Our Science in 2022 and currently serves on its Board of Directors.

### Data Sources

Publicly available information regarding stipends and waivers for the 38 universities in Canada offering graduate studies in Physics and/or Ecology (or the most similar department) was collected between February 16 to August 16, 2024. This study is limited to these two NSERC-funded programs to simplify the lengthy and difficult data collection process (see Accessibility), and because data are only valid for a single year. Tuition, mandatory fees, and minimum stipend amounts for each degree program were collected (see Supplementary Material for full details). When not discoverable on department websites or in associated files (e.g., graduate student handbooks), information was requested from program contacts via email.

### Accessibility

Determining accurate annual stipends was difficult, despite two of four authors possessing PhD’s. It was rare for tuition and fees to be displayed in logical and easily interpretable ways. For example, most institutions split fees into different categories (e.g., bus pass, graduate association fees), but they commonly use different units of time to display each: some fees appeared per term, some per year, some per credit, while others had different values during fall, winter, and summer or for part-time vs. full-time students, and so on. This complexity made it extremely easy to make simple errors due to the number of parameters and unclear language/presentation. Many institutions include certain fees (e.g., healthcare) on their fee lists, others do not. Furthermore, several institutions put dollar amounts behind several menus and/or ‘opt-out/opt-in’ paperwork, or even private intranet pages.

Whatever the factors contributing to low discoverability and transparency in tuition and stipend data may be, the impact is obfuscated financial information prior to being enrolled in a program. A solution for these issues would be to have a “Finances” page clearly indicated on department websites, with a table laying out common funding scenarios, minimum stipends, durations, costs, and how much of the stipend is left after fees and tuition. A department that models this well is the University of British Columbia Physics Department website (https://web.archive.org/web/20240501083537/https://phas.ubc.ca/graduate-program-financial-support).

### Stipend Amounts

We collected tuition, fee, and stipend data from public-facing program and/or university websites for both domestic and international MSc and PhD students. We focus our main analysis on domestic students due to the complexity and uncertainty around much of the international student data that was available. We include international data in the Supplementary Online Material for reference, but acknowledge the lower confidence in the provided values.

Data analysis was performed in R (v. 4.4.0^8^). All code and data are publicly archived on GitHub (https://github.com/UVicMicropaleo/Canadian-Minimum-Graduate-Stipends). We defined Gross Minimum Stipend (GMS) as the minimum annual stipend, Net Minimum Stipend (NMS) as the minimum annual stipend after both tuition and fees are repaid to the institution, and MBM Shortfall as NMS minus the inflation-adjusted Market Basket Measure poverty threshold (MBM), for a single individual with no dependents assessed by Statistics Canada^9^. The dataset contains 140 programs from 38 institutions. We found 91 programs providing a guaranteed minimum stipend, while 21 programs provide no minimum stipend. We could not identify minimums and received no response to email requests from 24 programs. Only considering departments which guarantee support, the mean domestic GMS in Canada is Can$23,878, and mean NMS is Can$16,621 (Figure 1). An average program with a minimum stipend charges Can$7,570 in tuition and fees to domestic graduate students, with a mean ∼34% (range: 18% - 61%) of the GMS repaid to their institution.

**Figure 1.**
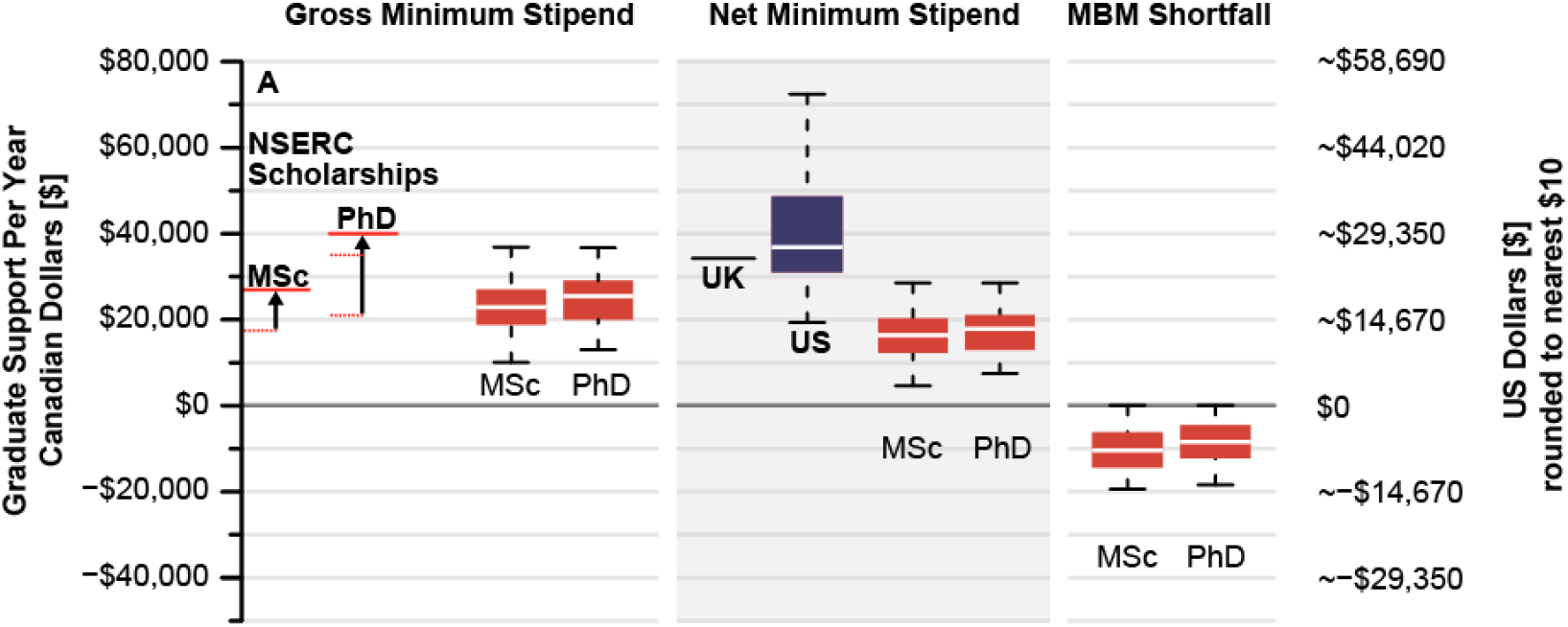
Annual financial support in Canadian and US dollars to graduate students (MSc and PhD in physics and biology). Red lines correspond to recently increased federal NSERC scholarships. Gross Minimum Stipend (GMS) is the guaranteed minimum funding for an MSc or PhD student provided by the institution. Net Minimum Stipend (NMS) is the GMS minus tuition and fees for an institution. MBM Shortfall is the NMS minus the Market Basket Measure (MBM), a poverty threshold, for an institution’s location. US Biology stipends are from ^3^ and should be considered a rough approximation of the US stipends, without accounting for fees or taxes (see text).

We compared stipend values to both governmental (MBM threshold^9^, figure 1) and private (a national report of rents for September 2023 by Rentals.ca^10^ both with and without adding $12k/yr for food and incidentals) metrics (Supplemental Information). Only two programs (University of Toronto Physics PhD and MSc) appear to have domestic net minimum stipends which reached MBM thresholds, though these were estimated as fees were hidden behind an intranet page. The average department would need to raise their guaranteed minimum stipend by ∼Can$9,468 for domestic students and ∼Can$16,899 for international students to reach the MBM threshold.

NMS best demonstrates the disparity between Canadian, US^11–13^, and UK stipends for PhD students (Figure 1a) as tuition waivers are normally included for US or UK graduate studies. Notably, US data^3^ (n=215) is considerable, but not exhaustive, while the UK funding agency sets a single national stipend level. It is difficult to reasonably compare various indices of poverty in an unbiased way across national borders. Poverty metrics generated by different governments vary due to numerous factors (e.g., national and regional differences, political impacts of declaring a ‘line of poverty’). Further, the non-TA portion of Canadian stipends are untaxed while US stipends are taxed. US stipend data from^3^ do not include ancillary fees, though it is extremely unlikely that fees or taxes are enough to erase the Can$23,666/yr difference between the means of the two countries. This difference is essentially the same as the mean GMS, suggesting in order to compete with the US or UK Canadian stipends would need to double, compared to only increasing by ∼1.5 times to meet the cost of living.

To identify if net stipend is driven by available institutional funding, we compared NMS against each University’s endowment (n = 26 universities with n=80 programs; Figure 2). We fit a generalized linear mixed model (package lme4 v.1.1.26,^14^) with university endowment (log10 transformed due to skewness of raw data), program (MSc or PhD), and field (Biology or Physics) as fixed effects, and university nested within province as a random effect because postsecondary funding falls within provincial jurisdiction. Endowment accounts for significant variation in NMS (Supplemental Table 2), suggesting a larger endowment generally results in higher stipends. Yet, substantial residual variation remains, even with accounting for variation across program and field within provinces (e.g., Ontario and Quebec demonstrate inter-institutional variation). Variation in stipends may result from a number of processes; e.g., different levels of support allocated to departments from the institution, different departmental budget priorities, or perhaps lagged responses to the increased expenses faced by students.

**Figure 2.**
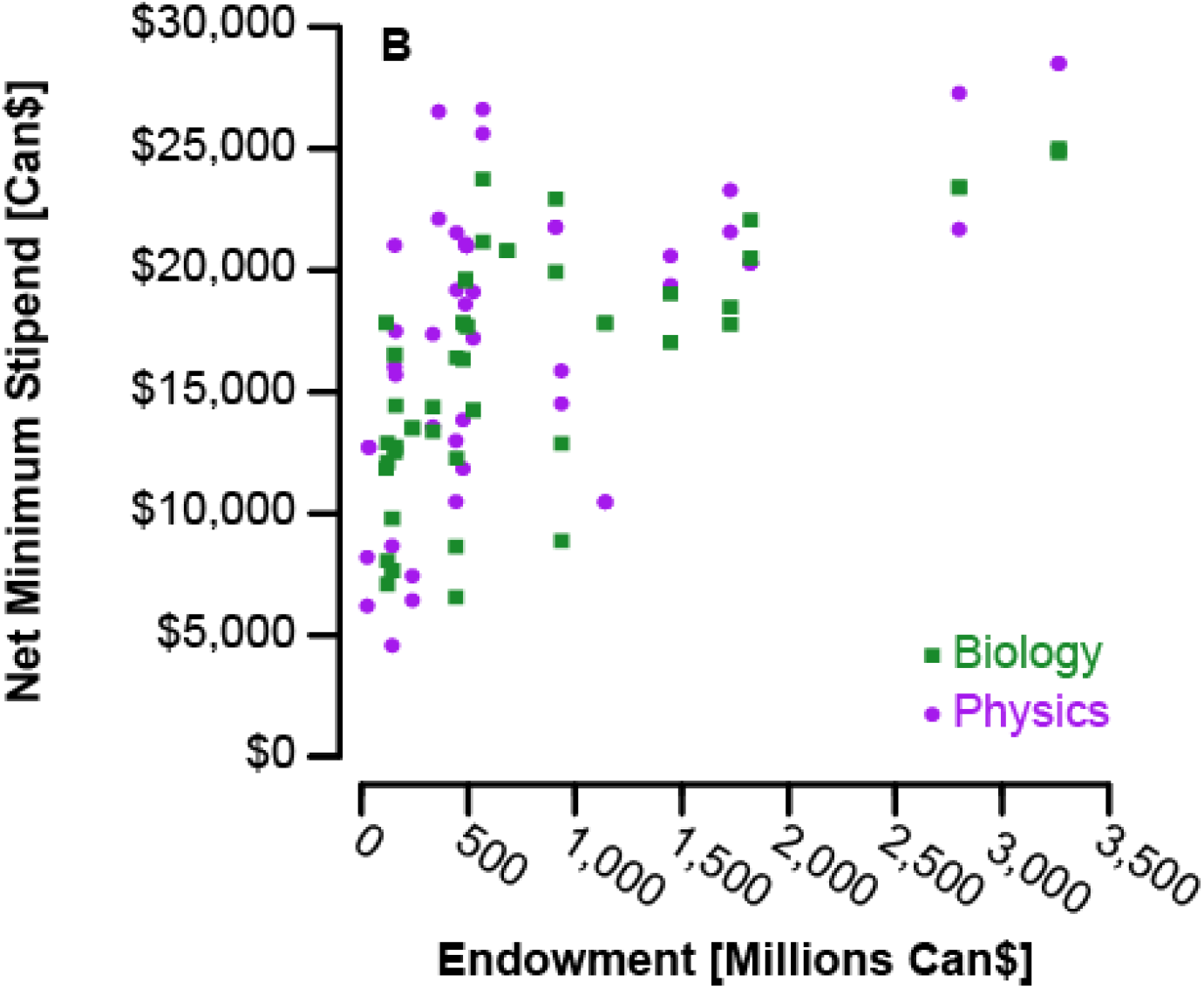
Net minimum stipends in Canadian dollars to MSc and PhD graduate students in biology (green squares) and physics (purple circles) in relation to their institution’s endowment in millions of Canadian dollars.

All analysis here is based on minimum stipend levels. Many graduate students do receive stipends higher than these levels, e.g. via top-ups from PI grants, additional TA hours, or external scholarships such as NSERC doctoral awards. However, the distribution of actual stipend values awarded is not publicly available and would require departments to report all stipend values, rather than simply department policies around guaranteed minimums (when they exist). Nevertheless, like minimum wage, minimum stipends are important to consider because they represent a lower bound that at least some graduate students experience. The purpose of setting a minimum is to ensure that all students receive financial support to enable them to focus on their studies without needing to seek employment elsewhere. If this is no longer being satisfied then the minimum level must be adjusted. Our mean/median minimum stipend levels are very similar to results from a recent survey^2^, suggesting many students are, or are close to, receiving their departments minimum stipend.

There has been significant advances for Canadian graduate students recently. Federally sponsored scholarships received a large increase in 2024^15^, giving all PhD student recipients positive net stipends after accounting for tuition and MBM thresholds. However, for the vast majority of graduate students in Canada who are financially supported through other means, minimum departmental stipends are still essential to ensure financial viability of graduate program enrollment. For example, in 2021-22 only ∼15% of domestic PhD students held a federal award. Further the proportion is likely to be lower for master’s students and near zero for international students who are ineligible for the majority of awards. Whilst these proportions will increase as more scholarships are awarded following Budget 2024, it will still remain the minority of students.

## Conclusions

The ‘hidden curriculum’ of graduate school admissions processes already results in attrition of individuals with identities underrepresented in the academy^16^. The complexity and opaque nature of the Canadian graduate funding system is a further barrier to recruiting underrepresented students, exacerbated by a lack of transparency around stipends and tuition at institutional and departmental levels. It also, given the dissimilarity with other similar nations, makes it incredibly challenging for international students to make informed decisions due to the overly-complex nature of tuition and fee structures. While individual departments are unable to transform this system, they can serve as the agents of change by improving transparency to allow for more informed choices by future graduate students and to guide applicants through understanding the reality of graduate research.

Canadian and foreign students at minimum stipend levels must incur substantial debt to undertake graduate educations, unless otherwise wealthy. This presents a series of problems for example: equity, the various health costs of poverty on the next generation of scientists, and the breadth of scientific inquiry. Students would benefit from significant changes at whatever level is possible^17^. Government, via funding agencies (e.g., NSERC;^18^), could effect the most change, with the ability to alter the entire system at once by enforcing a standard stipend level for all students, as seen in the UK and Australia, or requiring stipends to be tied to local cost of living metrics (like the MBM). However, within the current system, institutions, departments, or individual PIs can adjust minimum stipend levels. There is ample anecdotal evidence that this is already occurring at least at the departmental level, as several departments had increased stipends while we were performing quality control checks on our data. That stipends are still so low, however, suggests that individual departments cannot do this on their own, and require assistance from institutions and government.

It is clear that Canada is well behind competing countries when it comes to funding the next generation of scientists. Canadians who desire higher STEM education have three options: hope for significantly higher guaranteed support from a supervisor, department, or awards; incur substantial debts; or emigrate. If Canada still desires to attract and retain competitive, world-class talent, changes to Canadian graduate funding is required.

Graduate students, both in Canada and around the world, are the engine on which academic science runs^1^. Providing adequate financial support to these researchers enables this work to be done most effectively and boosts the ability of the entire scientific community to make progress. Not providing adequate support is damaging^19^. How best to provide this support is a global challenge, with countries offering a range of different systems and amounts.

However, regardless of the financial model offered, the productivity of science requires that we ensure graduate student researchers are not trapped under the poverty line.

## Acknowledgements

AJF is supported by a NSERC Discovery Grant (RGPIN-2022-03305) and ECR Launch supplement (DGECR-2022-00141). AEW, TB, and KK did not receive financial support for this research. M. Gaynor is thanked for their help with the US data. M. Gaynor, S. Laframboise, and K. Kharas are thanked for a preliminary review.

