## Supplemental Information for "Canadian science graduate stipends lie below the poverty line"

| **Name** | **Contributions** | **ORCiD** |
| --- | --- | --- |
| Thomas Bailey | Project administration, data management, data collection, analysis, writing | NA |
| Andy Fraass | Data collection, analysis, writing | 0000-0002-2007-3616 |
| Kayona Karunakumar | Data collection, writing | NA |
| Andrea Wishart | Data collection, analysis, writing | 0000-0001-7954-0613 |

**Affiliations**

Andrew Jeffrey Fraass^1,2,3†^ & Thomas J Bailey^4,5†^*, Kayona Karunakumar^4^, Andrea E. Wishart^4†^

Primary Affiliation: ^1^University of Victoria – School of Earth and Ocean Sciences, Bob Wright Centre A405, PO Box 1700 STN CSC, Victoria, B.C. V8W 2Y2 Canada, (250)721-6314,

^2^University of Bristol – School of Earth Sciences

^3^The Academy of Natural Sciences of Drexel University – Invertebrate Paleontology

^4^Support Our Science, Ottawa, ON, Canada

^5^Department of Physics, University of Ottawa, Ottawa, ON, Canada

, 343-262-6173

**Supplemental Information**

*Data collection*

In addition to tuition and stipend values as described in the main text, we also gathered the following information for each program: if funding was guaranteed and for how long, if waivers were provided to offset higher international tuition levels, and if teaching assistantship (TA) or research assistantships(RA) roles were required and if so, how many hours.

[Minimum Stipend Data](https://docs.google.com/spreadsheets/d/1STusmXjLqOZeInesHmFcUbIf_eWV9eDwFvfxobQ5LlE/edit?usp=sharing)

Data for this project can be found in multiple formats. A GitHub repository (<https://github.com/UVicMicropaleo/Canadian-Minimum-Graduate-Stipends>) hosts a machine readable file and all code associated with the project. Data collection was performed by sharing a GoogleSheet (<https://docs.google.com/spreadsheets/d/1STusmXjLqOZeInesHmFcUbIf_eWV9eDwFvfxobQ5LlE/edit?usp=sharing>) for all collaborators to use. In order to speed replication of this study across other fields or funding regimes, the authors invite others to mirror the spreadsheet and engage in their own data collection (in collaboration or not).

*Scoring data transparency for tuition and stipends*

Tuition Data was scored out of 4 for the level of ease it took to get to the selected program. Here, we worked with links through multiple web pages to obtain course-specific figures, assess how linked it was from the program site or through Google; and locate tuition values on the program site. This scoring system also assessed the level of ease to parse for the following: annual tuition located in one place (i.e., not only accessible by semester); annual fees clearly stated (i.e., if the program/student’s status only needed to be entered one time), and to what extent the information was shown on the website and not found in an external document.

Stipend Data was scored out of 6 for the level of ease it took to get to the web page which included stipend figures. This included assessing the following: if information was or was not available; if it was hard (i.e., if there was an undue amount of clicking required to get to the information); if it was moderate (i.e., consisting of more than two clicks from the program homepage or if the program homepage was slightly more challenging to find); or easy (i.e., consisting of one or two clicks from the program homepage or if the program homepage was slightly harder to locate). This scoring system also assessed to what extent complete information could be found regarding international students, the effect of scholarships, and the net stipend (after tuition).

| **Score item** | **Mean ± SD** |
| --- | --- |
| Tuition (out of 2) |  |
| Discoverability | - 0.7 ± 0.6 |
| Parse | - 1.2 ± 0.6 |
| Stipend (out of 3) |  |
| Discoverability | - 1.6 ± 1.3 |
| Complete | - 0.8 ± 1.1 |

**Supplemental Table 1.** Mean scores (± standard deviation) for discoverability and transparency of tuition and stipend data. Tuition transparency was assessed as ease of parsing, while stipend transparency was assessed as completeness of presented data. Scores were assigned based on the rubrics presented in Supplementary Online Material. Higher values correspond to greater discoverability and transparency, while lower values correspond to lower levels of both.

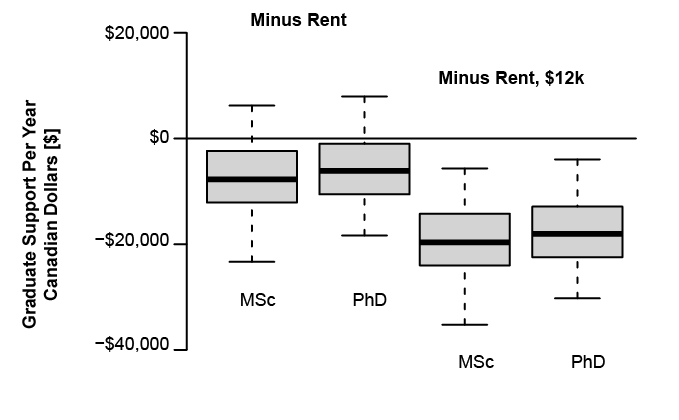

**Supplemental Figure 1.** Supported domestic minimum stipends compared to Rentals.ca values. Boxes at left are Net Minimum Stipends minus a local assessment of rents from Rentals.ca ^10^, while boxes at left subtract an additional $12k to account for food and other costs.

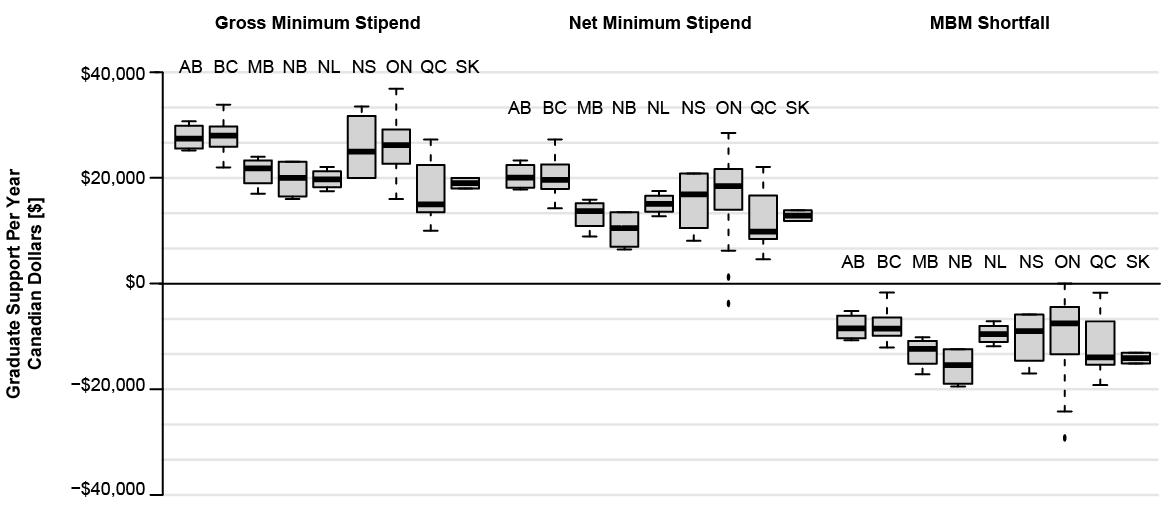

**Supplemental Figure 2.** Supported domestic minimum stipends divided by province. Gross Minimum Stipend (GMS) is the guaranteed funding for a student provided by the institution. Net Minimum Stipend (NMS) is the GMS minus tuition and fees for an institution. MBM Shortfall is the NMS minus the Market Basket Measure (MBM) for an institution's location.

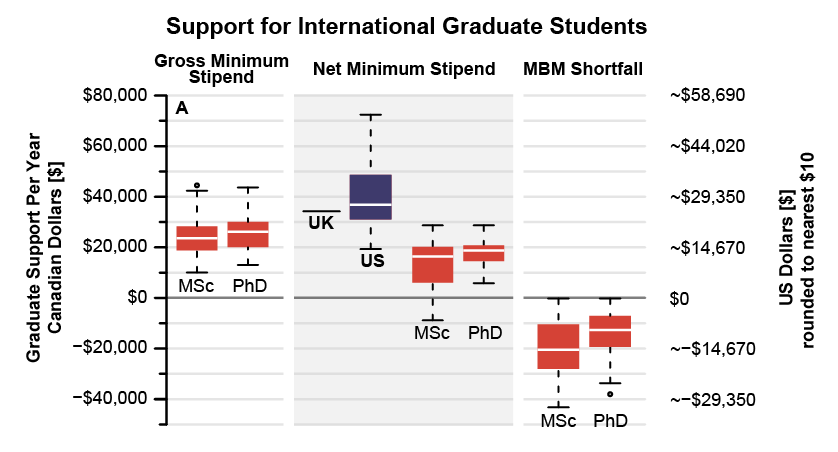

**Supplemental Figure 3.** Annual financial support in Canadian and US dollars to international graduate students (MSc and PhD in physics and biology). Gross Minimum Stipend (GMS) is the guaranteed minimum funding for an MSc or PhD student provided by the institution. Net Minimum Stipend (NMS) is the GMS minus tuition and fees for an institution. MBM Shortfall is the NMS minus the Market Basket Measure (MBM), a poverty threshold, for an institution's location. US Biology stipends are from ^3^ and should be considered a rough approximation of the US stipends, without accounting for fees or taxes (see text).

|  | **Estimate ± SEM** | **d.f.** | ***t*** | ***p*** |
| --- | --- | --- | --- | --- |
| **Fixed effects** |  |  |  |  |
| Intercept | -7447.1 ± 4055.6 | 22.99 | -1.84 | 0.079 |
| Endowment  (log_10_ transformed) | 8162.8 ± 1449.4 | 20.73 | 5.630 | < 0.001* |
| Program (reference group: MSc) | 1737.2 ± 523.5 | 523.5 | 52.81 | 0.002* |
| Field (reference group: Biology) | 849.2 ± 586.5 | 522.9 | 58.7 | 0.153 |
| **Random effects** | **Variance ± SD** |  |  |  |
| University:Province (Intercept) | 9352453 ± 3058 |  |  |  |
| Province (Intercept) | 3497893 ± 1870 |  |  |  |
| Residual | 5480617 ± 2341 |  |  |  |

**Supplemental Table 2.** Summary of generalized linear mixed model fit for Net Minimum Stipend (NMS) using package lme4 v. 1.1.26 (Bates et. al 2015). SEM = Standard error of the mean; SD = Standard deviation. Asterisk * denotes significance at α = 0.01.
